# Bioinformatics approach reveals Nur77-regulated genes common to the central nervous and immune systems

**DOI:** 10.1101/2020.05.28.120949

**Authors:** Montserrat Olivares-Costa, Guillermo Parada, María Estela Andrés

## Abstract

Neurons and immune cells have similar mechanisms for cell to cell communication. Exists active crosstalk between neurons and immune cells. Neurotransmitters can activate or inhibit immune cells, and cytokines modulate neuronal activity. The transcription factor Nur77 (NR4A1) is highly and transiently induced by several stimuli in neurons and T-lymphocytes, suggesting that Nur77 is involved in the coordination of the genetics response in both cell types. By analyzing publicly available databases, we found Nur77 target genes common to the nervous and immune systems. Functions as cell adhesion and anchoring appear as processes regulated by Nur77 in neurons and immune cells.

## Introduction

Nur77 (also known as NGFI-B, TR3, and NR4A1) [1,2,3] is a transcription factor encoded by an immediate-early gene that belongs to the nuclear receptor superfamily. Along with Nurr1 and Nor1 form a specific subgroup of orphan members, known as Nur subfamily. Crystallographic studies show that Nur77, like Nurr1, adopts a naturally active transcription conformation and lacks the canonical ligand-binding pocket present in ligand-inducible nuclear receptors [4,5]. All members of the Nur subfamily bind DNA as monomers through the Nerve-Growth-Factor Inducible gene B (NGFI-B)-responsive element (NBRE) (A/TAAAGGTCA) [6,7] and as homodimers or heterodimers to Nur-Responsive Element (NurRE) (AAATG/AC/TCA) [8]. Nur77 and Nurr1 can also form heterodimers with the retinoid x receptor (RXR) binding the genome on DR5 elements (GGTTCAnnnnnAGGTCA) [9].

Nur77 has been largely studied in the immune system for its function inducing apoptosis of auto-reactive immature lymphocytes T and modulating the inflammatory response [10,11]. In the Central Nervous System (CNS), Nur77 is widely expressed, particularly in brain nuclei that receive dopaminergic and noradrenergic neurotransmission, which regulates the expression of Nur77 [12]. Brain pathologies characterized by an imbalance of catecholamine neurotransmission, such as anxiety, addiction, and schizophrenia, are associated with changes in the expression of Nur77 [13,14]. Nur77 also seems to play a key role in neurodegenerative disorders. In models of Parkinson’s disease, Nur77 expression increases significantly in the substantia nigra, where it appears to be involved in the loss of dopamine cells since the genetic disruption of NR4A1 gene reduced the loss of dopamine neurons induced by neurotoxins [15].

A close relationship between the nervous and immune systems has been reported, existing not only physical contact through immune resident cells in the CNS, and nervous terminals that innervate peripheral immune organs [16] but also chemical communication. For instance, inflammatory cytokines modulate neuronal functions through receptors present in neurons and glial cells [17,18]. On the other hand, neurotransmitters as dopamine, glutamate, and serotonin released from neurons and immune cells, can activate or suppress T-cells in a context-dependent manner [19]. In turn, T-cells express receptors for many different neurotransmitters on their surface, and their expression is regulated upon time by T cell receptor (TCR) activation [19,20]. In addition, neurons and immune cells share several features like the capacity to establish connections and transmit information to neighboring cells through specialized cell-cell contacts, named synapsis in both systems [21].

In both the nervous and immune systems, Nur77 emerges as a key factor in the rapid signaling of cell-cell communication and the genetic plan that begins in the postsynaptic cell. In the CNS, the rapid and transient expression of Nur77 induced by neuronal activity regulates the distribution and density of dendritic spines in pyramidal neurons of the hippocampus [22], and the metabolism of neurons by regulating mitochondrial energetic competence under chronic stimulation [23]. In T cells, Nur77 is an activity marker downstream of TCR, the main receptor in immunological synapses [24,25]. Sekiya et al (2013) showed that strong TCR signals triggered by self-antigens induce Nur77 (and the other Nur factors), which in turn control the transcription of Foxp3, thus driving T regulatory developmental program [26]. Nur77 also modulates the inflammatory autoimmune response by regulating the metabolism and proliferation of T cells [27]. In addition, Nur77 regulates the production of norepinephrine in macrophages by inhibiting the expression of tyrosine hydroxylase, which limits experimental autoimmune encephalomyelitis [28,29].

Given that Nur77 is involved in the response to synaptic stimulation and control of metabolism in immune cells and neurons during their activation, we hypothesized that Nur77 regulates the expression of a set of genes common to the nervous and immune systems. To answer this question, we used bioinformatic approaches to find target genes of Nur77 common to both systems.

We used four different databases, publicly available. Our analysis reveals that there is a select group of common Nur77 target genes for the CNS and the immune system. The functions of cell adhesion and anchoring emerge as the processes regulated by Nur77 in both systems. We also characterized Nur77 binding sites throughout the genome, finding a significant distribution in strong enhancers and active promoters, reinforcing the function of Nur77 as a transcriptional activator in CNS and the immune system.

## Methods

### Data acquisition

We downloaded the ChIP-Seq peaks from Gene Expression Omnibus (GEO) corresponding to EGFP-Nur77 in K562 (GSE31363) [30] and endogenous Nur77 in NSC (GSM1603270) and NC (GSM1603273) [31]. The microarray data was taken from the work published by Chen et al. in 2014, (GEO Accession; GSE76805) [22]. We used ENSEMBL gene annotation for human (hg19) and mouse (mm10) which were directly extracted as TxDb objects though GenomicGeutres and TxDb. Mmusculus.UCSC.mm10.ensGene R packages. Additional genomic annotations of different chromatin states and functional regions were extracted from UCSC Table browser, including the K562 Genome Segmentation by ChromHMM and Ensembl Regulatory Build.

### Nur77 binding site characterization

The overlap between the different ChIP-Seq peaks and genomic features were calculated with the GenomicRanges R package using the function findOverlaps. We intersected the genomic coordinates with K562 Genome Segmentation by ChromHMM and calculated the overlapping enrichment, by normalizing by total coverage of each chromatin state across the genome.

To obtain a list of possible Nur77 target genes, we used the annotation of TSS and proximal promoters provided by the Ensembl Regulatory build for human (hg19) and mouse (mm10). The annotated TSS and proximal promoters were assigned to transcripts ID from Ensembl annotation if they were located within 2000 nt upstream and 500 nt downstream from a transcript start coordinate. Then, the function findOverlaps from GenomicRanges was used to find Nur77 ChIP-Seq peaks that overlapped with TSS or proximal promoters from the Ensembl Regulatory build. Finally, we used the biomaRt R package to find the common gene symbol associated with each of the Ensembl IDs that were assigned to each TSS and proximal promoter that overlapped with a Nur77 ChIP-Seq peak.

### Microarray analyses

We analyzed the gene expression changes after the overexpression of Nur77 in hippocampal pyramidal cells by analyzing publicly available microarray data published by Chen et al. in 2014 [22]. We computed the differentially expression results from the microarray data to filter genes that have an adjusted p-value lower than 0.05 and an absolute value of log2 fold change greater than 1. We plotted the results using ggplot2 and highlighting the names of genes for which a Nur77 peak were detected in their promoters.

### Ontology analysis

A list of common target genes of Nur77 in the neuronal and lymphatic systems was submitted to the Gene Ontology Consortium server (www.geneontology.org) [32,33] and processed using default parameters with PANTHER analysis tool [34].

## Results and discussions

### Characterization of Nur77 binding sites throughout the genome

To characterize Nur77 binding sites throughout the genome, we used a public database from the ENCODE project corresponding to a ChIP-Seq of overexpressing EGFP tagged Nur77 (EGFP-Nur77) in the *chronic myelogenous leukemia* cell line K562 (GEO accession: GSE31363) [30]. The analysis of the coordinates of EGFP-Nur77 peaks, reported by ENCODE, with respect to the nearest transcription start site (TSS), revealed a high frequency of Nur77 binding between +1000 and – 1000 nucleotides from TSS (Figures 1A, B).

**Figure 1:**
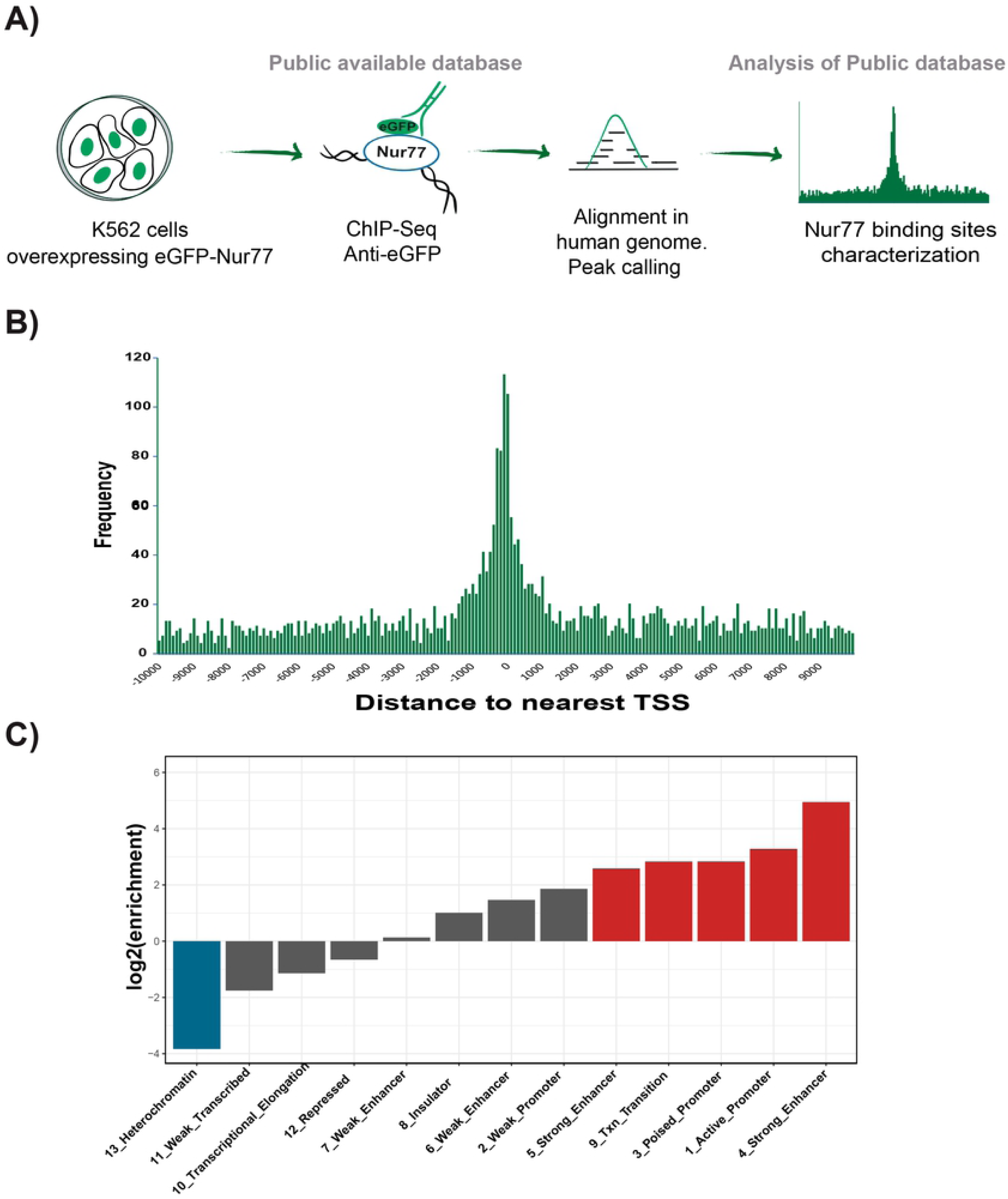
Nur77 binds to active promoters and strong enhancers in K562 cells. Analysis of publicly available database from the anti-EGFP ChIP-Seq experiment in K562 cell line overexpressing EGFP-Nur77 [30]. A) Pipeline of experimental procedure of public database and bioinformatics analysis. B) Abundance of Nur77 binding respecting to the nearest TSS location. C) Enrichment of Nur77 binding sites across different chromatin regions defined by Ernst et al. in 2011 [35]. In red, log2 of enrichment ≥ 2. In blue, log2 of enrichment ≤ -2. *

To further characterize the binding profile of EGFP-Nur77, we analyzed the enrichment of ChIP-Seq peaks across 15 chromatin states defined by Ernest et al. for K562 cell line [35]. These chromatin states (numbered from 1 to 15) were obtained through the implementation of multivariate hidden Markov models that integrates a wide range of epigenetic marks, including 8 histone posttranslational modifications, CTCF and H2A.Z histone variant [35]. We calculated the overlap enrichment across the EGFP-Nur77 peaks and chromatin states for K562 cell line, obtaining a significant enrichment of Nur77 peaks across chromatin states associated with transcription.

An enrichment analysis using the human chromatin segmentation model generated by Ernst, (2011) shows significant enrichment of Nur77 peaks in chromatin states associated with transcriptional regulatory regions, particularly in strong enhancers and active promoters (Fig. 1C). Two of chromatin states are described as strong enhancers (states 4 and 5), which differentiate in the occurrence of specific chromatin marks and distance to the TSS. Strong enhancer state number 4 presents a higher occurrence of histone 3 lysine 4 trimethylation (H3K4me3) and histone 3 lysine 9 acetylation (H3K9ac), and is closer to TSS than Strong enhancer state number 5. Nur77 is enriched in both chromatin states described as strong enhancers, exhibiting a log2 enrichment greater than 4 in chromatin state 4 (Fig. 1C). High enrichment of Nur77 binding is also observed in transcriptional transition states of chromatin. These areas present similar characteristics to transcriptional elongation areas, but with an increased presence of H4K20me1 and H3K4me1 and more sensitive to DNAse [35], suggesting an intermediate state between the promoter activation and effective elongation. In contrast, we found a negative enrichment for Nu77 binding in transcriptional elongation areas. Nur77 is poorly enriched in states numbers 6 and 7, both described as weak enhancers which differ in the occurrence of H3K4me2 and DNase sensitivity [35]. Finally, negative enrichment was observed in heterochromatin regions, indicating the absence of Nur77 in inactive areas of the chromatin (Fig. 1C).

Altogether these data indicate that Nur77 is mostly associated with active sites of chromatin, concordant with the role of Nur77 as a transcriptional activator, also confirmed by the high presence of Nur77 in the TSS. Our data also shows that Nur77 binds transcriptional transition areas, suggesting that Nur77 is present in the promoters of its target genes independently of their state (active or inactive). On the other hand, the data suggests that the presence of Nur77 in enhancers would be limited to the active state.

### Nur77 target genes in immune cells and neurons actively regulated by Nur77

We considered Nur77 target genes, those genes that bind Nur77 in their promoters (−2kb + 500 bp from TSS). To identify target genes of Nur77 in the CNS and the immune system, we analyzed published data sets. One set of data corresponds to the ChIP-Seq of overexpressed EGFP-Nur77 performed in K562 cell line [30]. Second and third data sets correspond to ChIP-Seqs of endogenous Nur77, of mouse neural stem cells (NSC) and NSC differentiated to neurons (NC) (GEO accession: GSM1603270 (NSC); GSM1603273 (NC)) [31]. From these sets of data, we identified the genes that bind Nur77 in their promoters. To learn what Nur77 target genes are actively regulated by Nur77, we crossed these data with the data of a microarray of mRNA from mouse hippocampal pyramidal neurons overexpressing Nur77 (GEO Accession; GSE76805) [22] (Fig. 2A).

**Figure 2:**
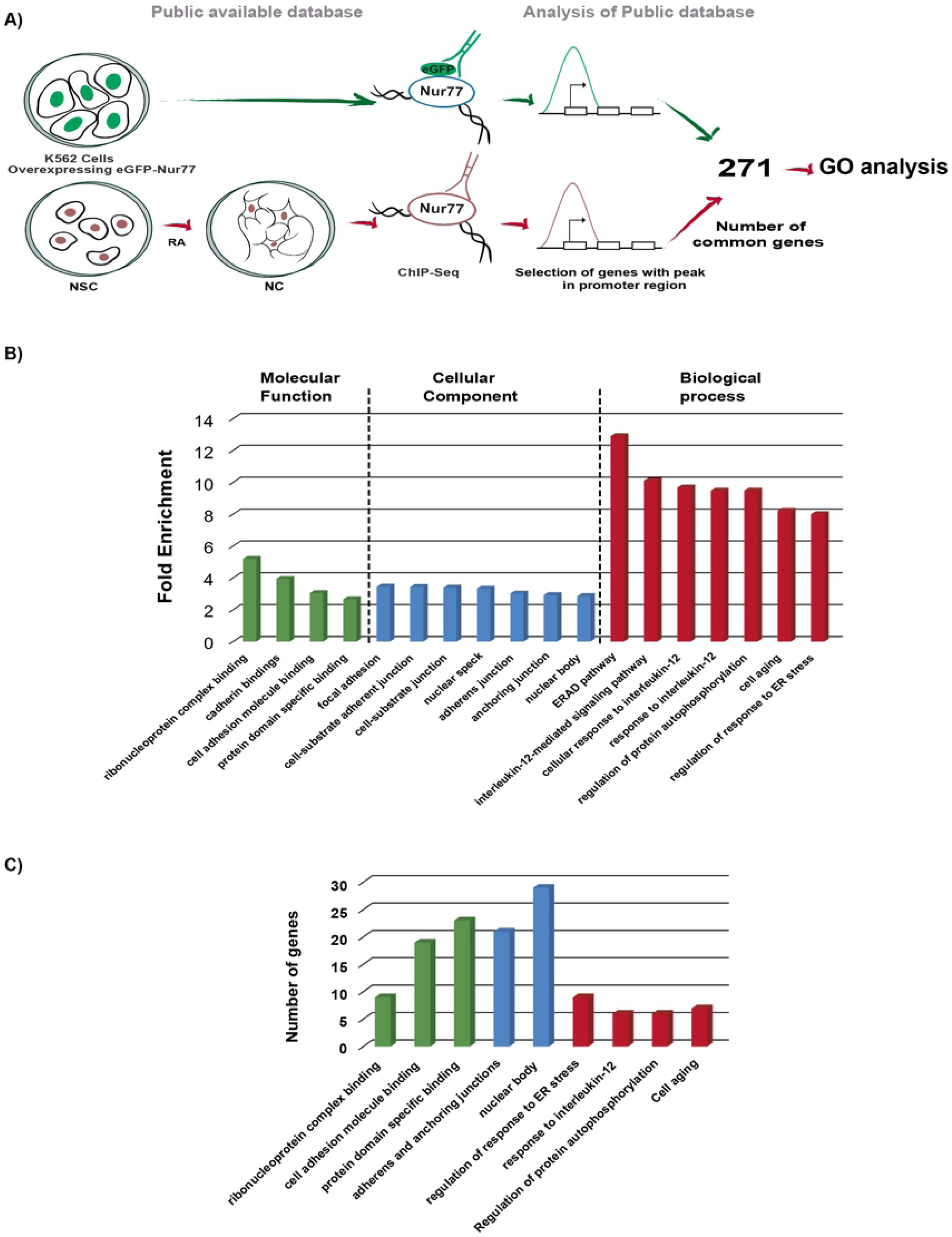
Nur77 target genes in neurons and immune cells. A) Pipeline of the experimental procedure of public databases [30,31] and the analysis to select Nur77 target genes. B,C,D) Volcano plots representing genes that change their expression when Nur77 is overexpressed in pyramidal neurons (blue), genes that also have a peak of binding of Nur77 in their promoter (red dots) and TSS (orange dots) according to the ChIP-Seq from K562 cells (B), ChIP-Seq form NSC (C) and ChIP-Seq from NC (D). Positive numbers indicate overexpression and negative numbers down-regulation. The dotted lines demarcate Log2 Fold of Change = ±1. *

The first analysis identified 58 genes that bind Nur77 in their promoters according to the ChIP-Seq of K562 cells, whose expression is modified in pyramidal neurons when Nur77 is overexpressed (Table S.1). Of these 58 genes, 9 changed their expression two or more times (Log2 fold of change ≥ 1) (identified with names in Fig. 2B). The second analysis identified 113 genes that bind Nur77 in their promoters according to the ChIP-Seq of NSC, whose expression is modified when Nur77 is overexpressed in pyramidal neurons (Table S.2). Of these 113 genes, 16 changed their expression two or more times (Log2 fold of change ≥ 1) (identified with names in Fig. 2C).

The third analysis identified 116 genes that bind Nur77 in their promoters according to the ChIP-Seq of NC, whose expression changed when Nur77 was overexpressed in pyramidal neurons (Table S.3). Of these 116 genes, 17 changed their expression two or more times (Log2 fold of change ≥ 1) (identified with names in Fig. 2D).

Although overexpression of EGFP-Nur77 in K562 cells resembles an active state, and therefore we cannot distinguish between the basal or induced attachment of Nur77, our analysis suggests that an important number of genes that bind Nur77 in their promoters are regulated by Nur77. This data also strengthens the idea that Nur77 is bound to the promoters of its target genes under basal conditions (NC and NSC), as it is shown in the previous analysis of the association of Nur77 to the different states of chromatin (Fig. 1C).

### Immune and nervous common Nur77 target genes

To find out whether Nur77 exerts a similar function in the nervous and immune systems, we searched for common target genes of Nur77 in neurons and immune cells. To this end, we selected genes that bind Nur77 on their promoters in both databases (1) the ChIP-Seq of K562 overexpressing EGFP-Nur77 [30] and (2) the ChIP-Seq of NC [31] (Fig. 3A). We found 271 target genes of Nur77 common to immune and neuronal ChIP-Seqs (Table S.4) (Fig. 3A). We are aware that subtle differences in ChIP-Seq protocols [30,31] could limit the common data when crossing both databases. However, we selected these databases given their high quality.

**Figure 3:**
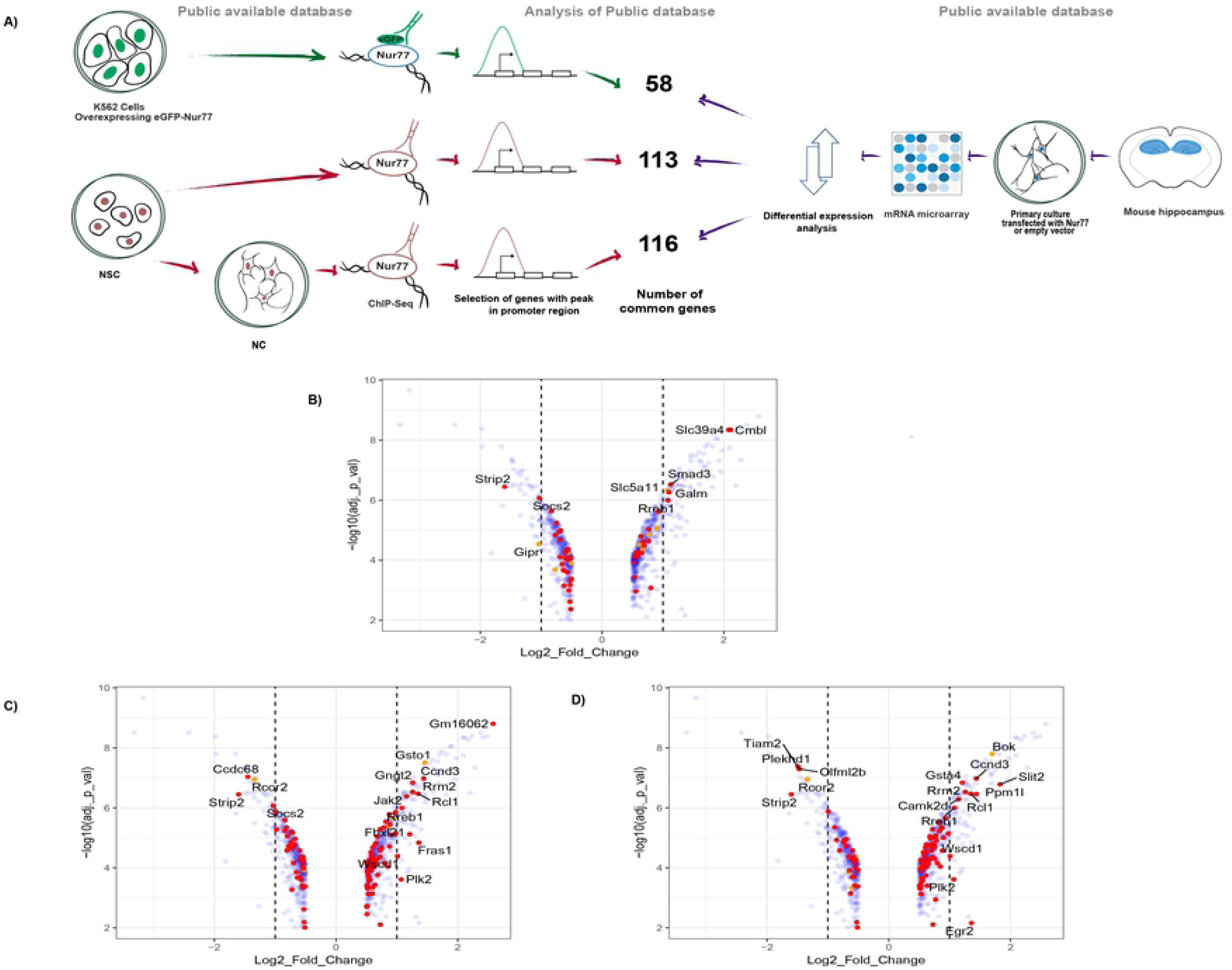
GO analysis of common target genes for Nur77 among the immune and nervous systems. A) Pipeline of experimental procedure of public databases, and corresponding GO analysis. B) Enriched GO terms (more than 2.5 fold of change). C) The number of genes classified in each GO terms category. Molecular function classification in green bars, cellular component classification in blue bars and biological process classification in red bars. Cell adhesion molecule binding category includes cadherin binding GO term genes. Adherens and anchoring junctions category includes genes of adherens junction, anchoring junction, focal adhesion, cell-substrate adherent junction, and cell-substrate junction GO terms. Nuclear body category includes genes of nuclear speck GO term. Regulation of response to endoplasmic reticulum stress includes genes of regulation of ERAD pathway GO term. Response to interleukin-12 category includes genes of interleukin-12-mediated signaling pathways, cellular response to interleukin-12 and response to interleukin-12 GO terms (Table S.5). *

We carried out Gene Ontology (GO) analysis of these genes for the three principal classifications that are molecular function, cellular components and biological processes (Fig. 3B and Table S.5). GO analysis showed an enrichment of Nur77 target genes in binding functions of the molecular function classification, which are: ribonucleoprotein complex binding (fold of enrichment 5.2), cadherin bindings (fold of enrichment 3.94), cell adhesion molecule binding (fold of enrichment 3.05) and protein domain specific binding (fold of enrichment 2.66) (Fig. 3B). In the cellular component classification, Nur77 target genes are enriched principally in categories of adhesion and junction: focal adhesion (fold of enrichment 3.45), cell-substrate adherent junction (fold of enrichment 3.43), cell-substrate junction (fold of enrichment 3.39), adherens junction (fold of enrichment 3.02), and anchoring junction (fold of enrichment 2.93). Two categories of nuclear localization were enriched too: nuclear speck (fold of enrichment 3.33) and nuclear body (fold of enrichment 2.86) (Fig. 3B). Many proteins encoded by Nur77 target genes, which are common to the nervous and immune system, are ribonucleoproteins and adhesion molecules. This fact is strengthened by the enrichment of Nur77 target genes in nuclear bodies and areas of adherents and anchoring junctions (Fig. 3C).

In the biological process classification, Nur77 target genes are enriched in regulation of Endoplasmic-Reticulum-Associated protein Degradation (ERAD) pathway (fold of enrichment 12.91) and regulation of response to endoplasmic reticulum stress (fold of enrichment 8.01). Three GO terms related to interleukin signaling were enriched: interleukin-12-mediated signaling pathway (fold of enrichment 10.11) cellular response to interleukin-12 (fold of enrichment 9.68) and response to interleukin-12 (fold of enrichment 9.49). Nur77 target genes were also enriched in the regulation of protein autophosphorylation (fold of enrichment 9.49) and cell aging (8.22) GO terms (Fig. 3B).

Crossing the database of genes that bind Nur77 in its promoters in both the immune and nervous systems (271 genes), with the database of the genes that are modified by increasing Nur77 in pyramidal neurons [22], it gives a set of 9 genes: AGAP3, BIRC5, DYM, ITGB3, KIF21B, MORN5, RREB1, STRIP2 and WEE1. Previous evidence supports that Nur77 controls the expression of BIRC5 gene [36], validating our results. Further studies are required to fully validate Nur77’s control over these genes, both in the nervous and immune systems.

In conclusion, our data analyses show that Nur77 is bound to the promoter of its target genes independently of their transcriptional state (weak, poised or active). In addition, our data suggest that the presence of Nur77 in enhancers is limited to the active state. We propose that Nur77 is always present in promoters but only binds to enhancers when it is upregulated, modulating thus transcription in response to stimuli.

Our analysis showed a strong participation of Nur77 target genes in anchoring and adhesion functions, which is consistent with the previously described roles of Nur77 in the modulation of neurite growth in neurons [23], and in the immune response [24,25], both processes that require interaction between cells and with the extracellular matrix.

Finally, the genes found in this work as common targets of Nur77 in the nervous and immune systems are new, undescribed targets of this transcription factor. The work presented here is an approach pretending to guide the experimental focus regarding Nur77 investigation, solving in part the problem of lack of knowledge of Nur77 target genes and presenting new functions that can be attributed to this transcription factor in both the immune and nervous systems.

## Supporting information

**Table S.1: Nur77 target genes in the K562 cell line**.

Genes that bind Nur77 on their promoter regions, according to the ChIP-Seq of K562 cells, and also modified their expressions when Nur77 is overexpressed in pyramidal neurons. Columns correspond to gene name, gene description from wikigene database [37], Nur77 peak location according to the ChIP-Seq from K562 cells (GSE31363) [30], Log2 of the fold of change and p-value adjusted according to differential expression analysis of microarray from pyramidal neurons overexpressing Nur77 (GSE76805) [22] and gene identifier in the ensembl database.

**Table S.2: Nur77 target genes in NSC**.

Genes that bind Nur77 in their promoter regions, according to the ChIP-Seq of NSC, and also modified their expressions when Nur77 is overexpressed in pyramidal neurons. Columns correspond to gene name, gene description from wikigene database [37], Nur77 peak location according to the ChIP-Seq from NSC (GSM1603270) [31], Log2 of the fold of change and p-value adjusted according to differential expression analysis of microarray from pyramidal neurons overexpressing Nur77 (GSE76805) [22] and gene identifier in the ensembl database.

**Table S.3: Nur77 target genes in NC**.

List of genes that bind Nur77 in their promoter regions, according to the ChIP-Seq of NC, and also modified their expressions when Nur77 is overexpressed in pyramidal neurons. Columns correspond to gene name, gene description from wikigene database [37], Nur77 peak location according to the ChIP-Seq from NC (GSM1603273) [31], Log2 of the fold of change and p-value adjusted according to differential expression analysis of microarray from pyramidal neurons overexpressing Nur77 (GSE76805) [22] and gene identifier in the ensembl database.

**Table S.4: Lymphatic and neuronal common target genes**

Target genes of Nur77 common to immune and neuronal systems. Are listed only the genes that present Nur77 binding on their promoter region. This set of genes was used to GO analysis.

**Table S.5: GO terms and genes classified on each one**.

GO terms enriched in the GO analysis and Nur77 target genes classified in each term. One table to each of the three principals GO categories are shown in separate sheets.

